# Circulating plasma IL-13 and periostin are dysregulated type 2 inflammatory biomarkers in prurigo nodularis: a cluster analysis

**DOI:** 10.1101/2022.06.07.495051

**Authors:** Varsha Parthasarathy, Karen Cravero, Junwen Deng, Zhe Sun, Sarah Engle, Autum Auxier, Nathan Hahn, Jonathan T. Sims, Angela Okragly, Martin P. Alphonse, Shawn G. Kwatra

## Abstract

**Background:** Prurigo nodularis (PN) is a chronic inflammatory skin disease characterized by severe pruritus and notable disease heterogeneity. There is evidence of systemic inflammation in PN, including dysregulation observed in type 2 inflammation in subsets of patients. We aimed to elucidate which components of type 2 inflammation are dysregulated in PN patients using plasma immunoassay cytokine profiling.

**Materials and Methods:** Whole blood was obtained from PN patients with uncontrolled disease and control patients without pruritus. Plasma samples were isolated from whole blood and assayed for IL-4, IL-5, IL-13, IgE, and periostin. For statistical analysis, ANOVA was utilized to compare PN and control patients. For multiple hypothesis, adjusted p-value was calculated with a Benjamini-Hochberg procedure with the significance threshold at 0.05. Clustering was performed using K-means clustering in R version 4.0.3.

**Results:** Single-plex assays of the T_h_2 biomarkers demonstrated significantly elevated circulating plasma IL-13 (0.13 vs. 0.006 pg/mL, p=0.0008) and periostin (80.3 vs. 60.2 ng/mL, p=0.012) in PN patients compared to controls. IL-4 (0.11 vs. 0.02 pg/mL, p=0.30) and IL-5 (0.75 vs. 0.40 pg/mL, p=0.10) were not significantly elevated, while IgE approached significance in PN (1202.0 vs. 432.7 ng/mL, p=0.08). Clustering of PN and control patients together revealed mean silhouette maximization at two clusters. Cluster 1 (n=36) consisted of 18 PN patients and 18 controls. Cluster 2 (n=11) consisted entirely of PN patients (p<0.01) (Figure 1b). When comparing the two clusters, cluster 2 had higher levels of IL-13 (0.33 vs. 0.008 pg/mL, p=0.0001) and IL-5 (1.22 vs. 0.43 pg/mL, p=0.03) compared to cluster 1. There were no significant differences in any of the biomarkers between PN patients from cluster 1 and the healthy control patients in this cluster.

Figure 1.
Single-plex plasma immunoassays of prurigo nodularis vs. controls (a) Plasma single-plex immunoassays (b) Heatmap of Z-scored biomarker levels for each patient, delineated by cluster. *PN, prurigo nodularis; HC, healthy control*

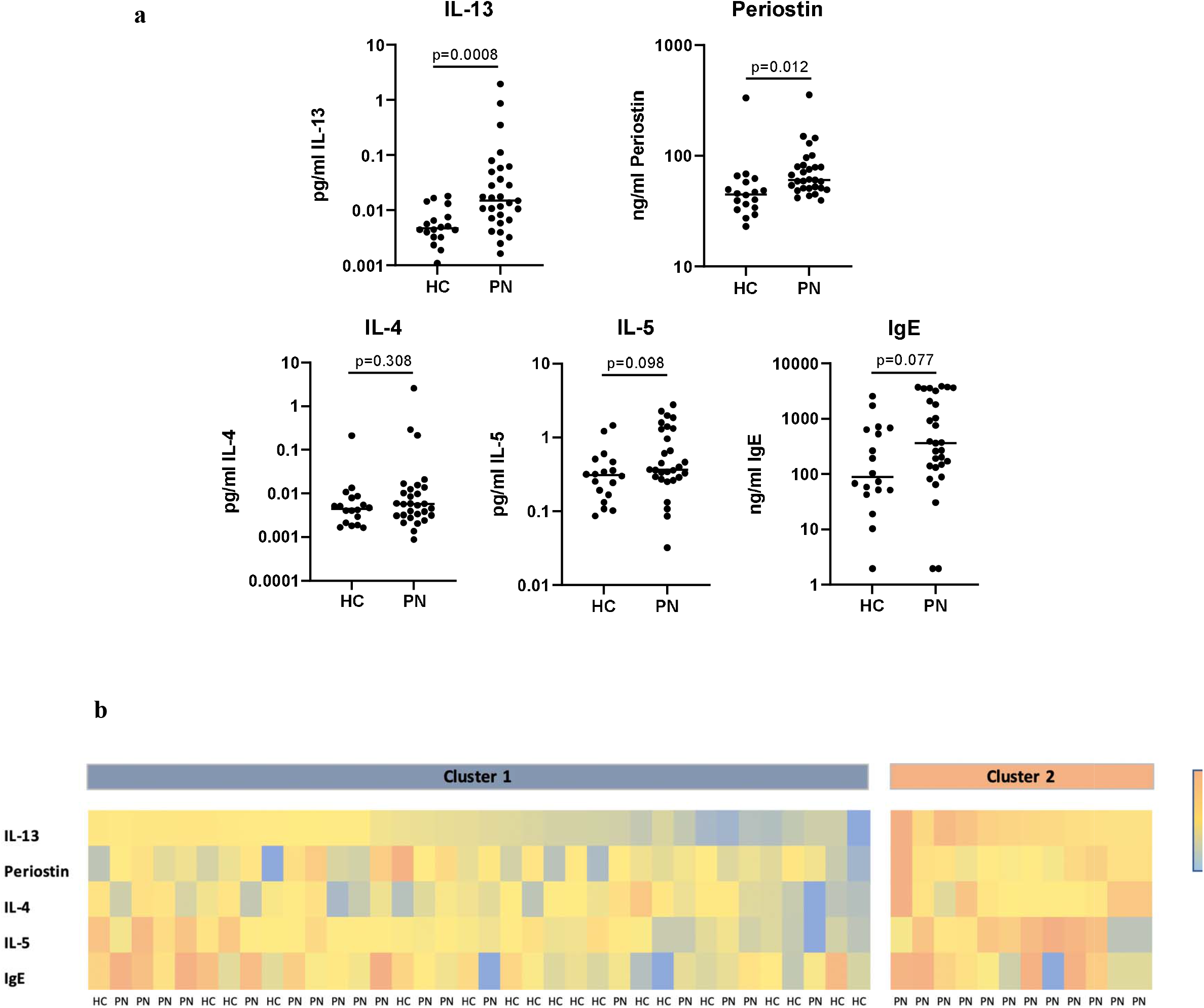

**Conclusions:** This study demonstrates elevation of IL-13 and periostin in PN patients with distinct clusters with varying degrees of type 2 inflammation. Given this heterogeneity, further biomarker studies with single-cell resolution will provide better guidance towards future precision medicine approaches in the treatment of PN.

---

Prurigo nodularis (PN) is a chronic heterogeneous inflammatory skin disease that presents with extremely itchy, hyperkeratotic nodules on the extremities and trunk, dramatically reducing quality of life for patients.^1^ Recent studies have identified increased CD4^+^, CD8^+^, γδ, and natural killer T cells in circulating peripheral blood mononuclear cells (PBMCs), suggesting PN is associated with systemic inflammation.^2^ There is also evidence of T_h_1, T_h_2, T_h_17, and T_h_22 involvement in PN in both skin and blood.^2,3^ The role of type 2 inflammation is of special importance in PN, given that there have been several off-label reports of the efficacy of treatment response with dupilumab, an IL-4 receptor α inhibitor that prevents IL-4 and IL-13 signaling.^4^ However, PN is unique from atopic dermatitis, a classical type 2 inflammatory disease, because it features a novel clinical presentation and unique neurovascular and fibrotic gene dysregulation.^5^ Therefore, we aimed to elucidate which components of type 2 inflammation are dysregulated in PN patients using plasma immunoassay cytokine profiling.

Whole blood was obtained from PN patients with uncontrolled disease and control patients without pruritus under a Johns Hopkins IRB-approved study (IRB00231694). Plasma samples were isolated from whole blood and assayed for IL-4, IL-5, IL-13, IgE, and periostin. IL-4 and IL-5 were measured with S-PLEX Human IL-4 and IL-5 kits (MSD, Rockville, MD). IL-13 was measured using the Simoa® human IL-13 Advantage HD-1/HD-X kit (Quanterix, Billerica, MA). IgE was measured with Invitrogen human IgE ELISA kit and periostin was assayed with the Invitrogen human periostin ELISA kit (ThermoFisher Scientific, Frederick, MD). For statistical analysis, ANOVA was utilized to compare PN and control patients using log-transformed data. For between-markers multiplicity adjustment, adjusted p-value was calculated with a Benjamini-Hochberg procedure with the significance threshold at 0.05. Clustering of PN and control patients was performed using K-means clustering in R version 4.0.3.

PN patients (n=29) and controls (n=18) had similar age (53.8 ± 13.4 vs. 52.2 ± 13.2 years, p=0.68), sex (72% vs. 72% female, p=0.98), and race (21% vs. 33% Caucasian and 69% vs. 67% African American, p=0.46) distributions. PN patients had higher Worst Itch Numeric Rating Scale scores (8.9 ± 1.3 vs. 0 ± 0) and Investigator Global Assessment scale scores (3.3 ± 0.6 vs. 0 ± 0) compared to controls.

Single-plex assays of the T_h_2 biomarkers demonstrated significantly elevated circulating plasma IL-13 (0.13 vs. 0.006 pg/mL, p=0.0008) and periostin (80.3 vs. 60.2 ng/mL, p=0.012) in PN patients compared to controls. IL-4 (0.11 vs. 0.02 pg/mL, p=0.30) and IL-5 (0.75 vs. 0.40 pg/mL, p=0.10) were not significantly elevated, while IgE approached significance in PN (1202.0 vs. 432.7 ng/mL, p=0.08) (Figure 1a).

Clustering of PN and control patients together revealed mean silhouette maximization at two clusters. Cluster 1 (n=36) consisted of 18 PN patients and 18 controls. Cluster 2 (n=11) consisted entirely of PN patients (p<0.01) (Figure 1b). When comparing the two clusters, cluster 2 had higher levels of IL-13 (0.33 vs. 0.008 pg/mL, p=0.0001) and IL-5 (1.22 vs. 0.43 pg/mL, p=0.03) compared to cluster 1. There were no differences in periostin (p=0.06), IL-4 (p=0.07), or IgE (p=0.41) between these groups. PN patients from cluster 2 (n=11) demonstrated higher levels of IL-13 than cluster 1 PN patients (0.33 pg/mL in cluster 2 vs. 0.007 pg/mL in cluster 1, p=0.0003). There were no significant differences in any of the biomarkers between PN patients from cluster 1 and the healthy control patients in this cluster.

IL-13 and periostin both stimulate the release of IL-31 from inflammatory cells, including macrophages and eosinophils, propagating systemic inflammation in PN.^6^ Previous studies have also found that there are endotypes in PN, represented by varying degrees of circulating inflammatory biomarkers across varying immune axes, including IL-1α, IL-4, IL-5, IL-6, IL-10, IL-17A, IL-22, IL-25, and IFN-α.^7,8^ Our findings support distinct endotypes in PN with respect to type 2 inflammation, with one cluster consisting solely of PN patients with elevated IL-13 compared to another less inflammatory cluster, which showed no significant differences between PN and control patients with respect to T_h_2 biomarkers. This supports clinical observations of heterogeneity in response to T_h_2-modulating therapies in PN patients.^4^ Limitations of this study include a cross-sectional design which may prevent inference of causation. In summary, this study demonstrates elevation of IL-13 and periostin in PN patients with distinct clusters with varying degrees of type 2 inflammation. Given this heterogeneity, further biomarker studies with single-cell resolution will provide better guidance towards future precision medicine approaches in the treatment of PN.

